# *bamSliceR*: a Bioconductor package for rapid, cross-cohort variant and allelic bias analysis

**DOI:** 10.1101/2023.09.15.558026

**Authors:** Yizhou Peter Huang, Lauren Harmon, Eve Deering-Gardner, Xiaotu Ma, Josiah Harsh, Zhaoyu Xue, Hong Wen, Marcel Ramos, Sean Davis, Timothy J. Triche

## Abstract

The NCI Genomic Data Commons (GDC) provides controlled access to sequencing data from thousands of subjects, enabling large-scale study of impactful genetic alterations such as simple and complex germline and structural variants. However, efficient analysis requires significant computational resources and expertise, especially when recalling variants from raw sequence reads. We thus developed ***bamSliceR***, an R/Bioconductor package that builds upon the ***GenomicDataCommons*** package to extract aligned sequence reads from cross-GDC meta-cohorts, followed by targeted analysis of variants and effects (including transcript-aware variant annotation from transcriptome-aligned GDC RNA data). Here we demonstrate population-scale genomic & transcriptomic analyses with minimal compute burden via ***bamSliceR***, identifying recurrent, clinically relevant sequence and structural variants in the TARGET AML and BEAT-AML cohorts. We then validate results in the (non-GDC) Leucegene cohort, demonstrating how the ***bamSliceR*** pipeline can be seamlessly applied to replicate findings in non-GDC cohorts. These variants directly yield clinically impactful and biologically testable hypotheses for mechanistic investigation. ***bamSliceR*** has been submitted to the Bioconductor project, where it is presently under review, and is available on GitHub at https://github.com/trichelab/bamSliceR.

## 1 INTRODUCTION

Detection of genetic variants from whole exome sequencing (WXS) or whole genome sequencing (WGS), has unveiled novel mechanisms in rare diseases and cancers. The consequences of these variants can often be further illuminated by whole-transcriptome sequencing (WTS) via RNA sequencing (RNA-seq). When discrepancies are observed between genomic and transcriptomic dosage of variant allele frequencies (VAFs), commonly referred to as *allelic bias*, this can offer important insights into the mechanisms of disease initiation, progression, and response to therapy. Furthermore, the predicted consequences of transcribed variants depend upon the specific transcripts in which they appear. Combining WGS/WXS with WTS can provide direct evidence for the most relevant transcript or transcripts impacted by allelic variation, which can be particularly important when variants of interest primarily occur in rare or novel transcripts. Recurrent allelic bias affects *TP53*- and *WT1*-mutant disease, and perhaps most notably, *MECOM*-rearranged leukemia (Gröschel et al. 2014), which exclusively activates the oncogenic short transcript encoding the EVI1 proteoform. Allelic bias can also buffer the impact of variants in genes such as the X-linked *KDM5C* locus, where skewed X chromosome inactivation appears to protect female carriers from severe neurodevelopmental and immune phenotypes observed in male variant carriers (Giovenino et al. 2023). Somatic loss of heterozygosity is a primary mechanism of progression for germline risk variants such as those seen in *BRCA1*(Cancer Genome Atlas Research Network), where promoter DNA methylation leads to monoallelic expression of only the variant allele.

However, logistical and financial constraints can preclude generation of paired genomic and transcriptomic data, and a recent surge in RNA-seq data from clinical trials has surpassed WGS as a primary tool to assess expressed genomic variants (Casamassimi, *et al*. 2017). For example, the Leucegene project generated publicly available RNA-seq data from acute myeloid leukemia (AML) patients (n=452) and developed a local assembly approach to identify mutations in raw RNA-seq reads (Audemard, *et al*. 2019). While this method is computationally efficient, it involves re-alignment and does not take advantage of key features in the Binary Alignment and Mapping (BAM) (Li, *et al*. 2009) format. Large consortia, including The Cancer Genome Atlas (Cancer Genmoe Atlas Research Network 2013) (TCGA), TARGET (Bolouri, *et al*. 2018), and BEAT-AML (Tyner, *et al*. 2018), collected RNA-seq data as a biological correlative at rates outpacing WGS in the same cohorts and studies. The NCI GDC provides controlled access to harmonized clinical and genomic data from these and other consortia with a high-performance infrastructure (Wilson, *et al*. 2017). Our package takes advantage of the GDC-managed BAM-slicing API to automate inspection of specific genome regions across large GDC cohorts using BAM indices (Li, *et al*. 2009; Wilson, *et al*. 2017), providing an efficient option to fetch aligned reads for variant analysis.

Our R/Bioconductor software package, ***bamSliceR***, has been optimized to perform rapid variant characterization across thousands of patients’ sequencing data via GDC, SRA, or local files. ***bamSliceR*** takes advantage of GDC’s standardized BAM format and slicing API to efficiently extract reads from BAM files based on user-defined genomic ranges. Then, ***bamSliceR*** provides downstream functionality to facilitate tally read counts, annotation, and effect predictions of variants (Figure 1A). While the GDC BAM-slicing API does not support slicing transcriptome-aligned BAM files, we show that this can be accomplished via syntax familiar from *samtools*. Targeted transcriptome BAM files can also be generated from sliced, genomic BAM files by realigning against a transcript reference, as via *minimap2* (Li, *et al*. 2018) (Figure 1A). Finally, ***bamSliceR*** reports read counts for the reference and variant allele depths, the VAF estimates across experiments and alignment workflows, in many cases yielding genomic DNA VAF, genomic RNA VAF, and transcript-wise RNA VAF. These metrics are essential in modeling the consequence of variants linked to disease-related perturbations (Figure 1A).

**Figure 1.**
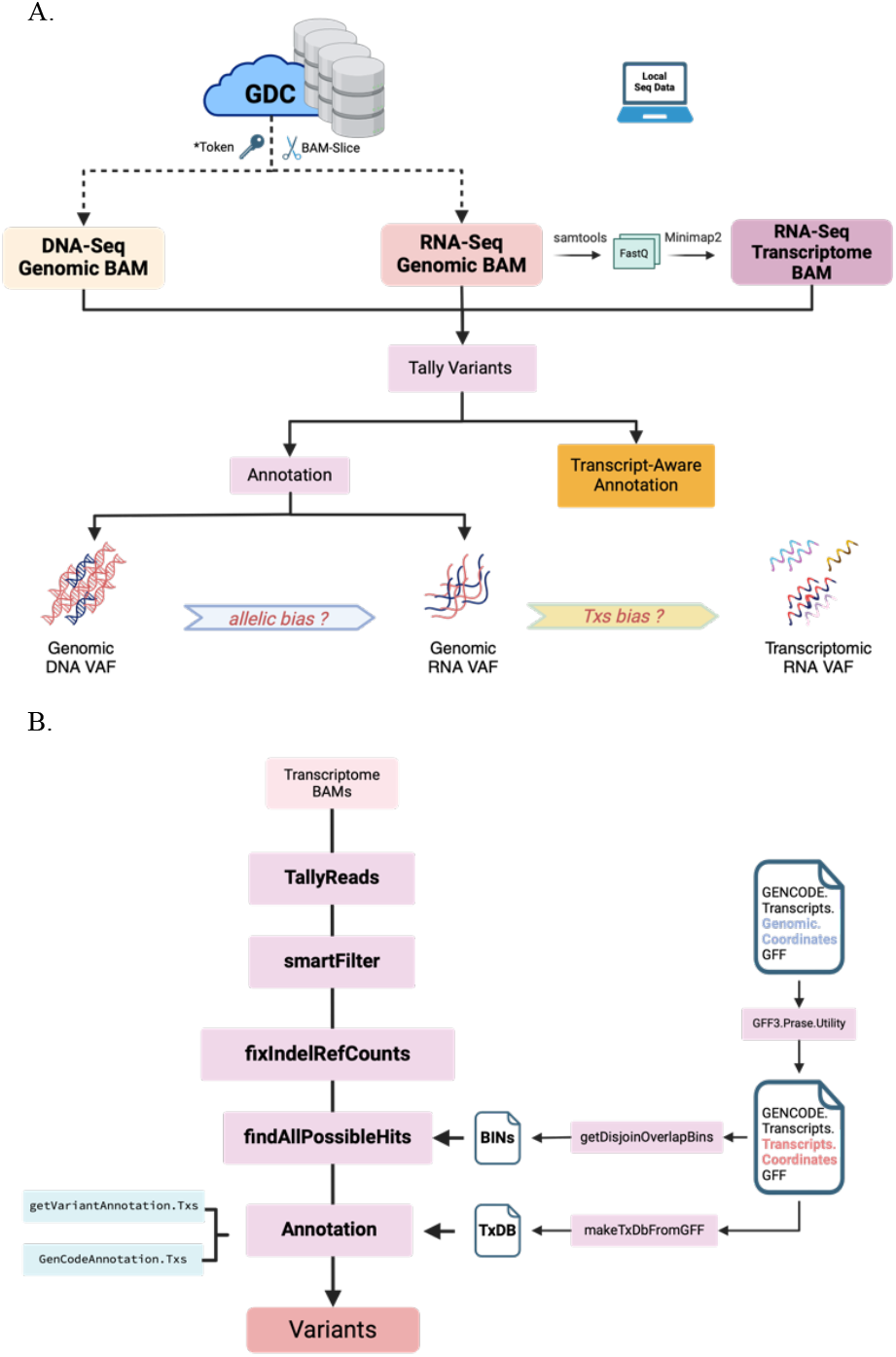
A) Overview of the *bamSliceR* Scheme. A. This schematic illustrates the bamSliceR pipeline, designed to efficiently retrieve metrics of variants from target regions across various data types, including DNA-Seq genomic BAMs, RNA-Seq genomic BAMs, and RNA-Seq transcriptome BAMs. **B) *bamSliceR* utility to retrieve metrics of variants fromRNA-Seq transcriptome BAMs**.

Here, we analyzed DNA-seq and RNA-seq data from more than 3,000 patients to demonstrate how our lightweight workflow can quickly investigate novel genomic regions of interest, increasing the scale and scope for discovery and validation. We demonstrate that ***bamSliceR*** bypasses the historically tedious variant calling workflow, reducing time, space, and computational burden by orders of magnitude relative to standard methods. With ***bamSliceR***, users can quickly transition from raw alignment data to variant characterization and perform population-scale targeted genomic data analysis.

## 2 Package Functionality

***bamSliceR*** has four main functions, as illustrated in Figure 2, and described here.

**Figure 2.**
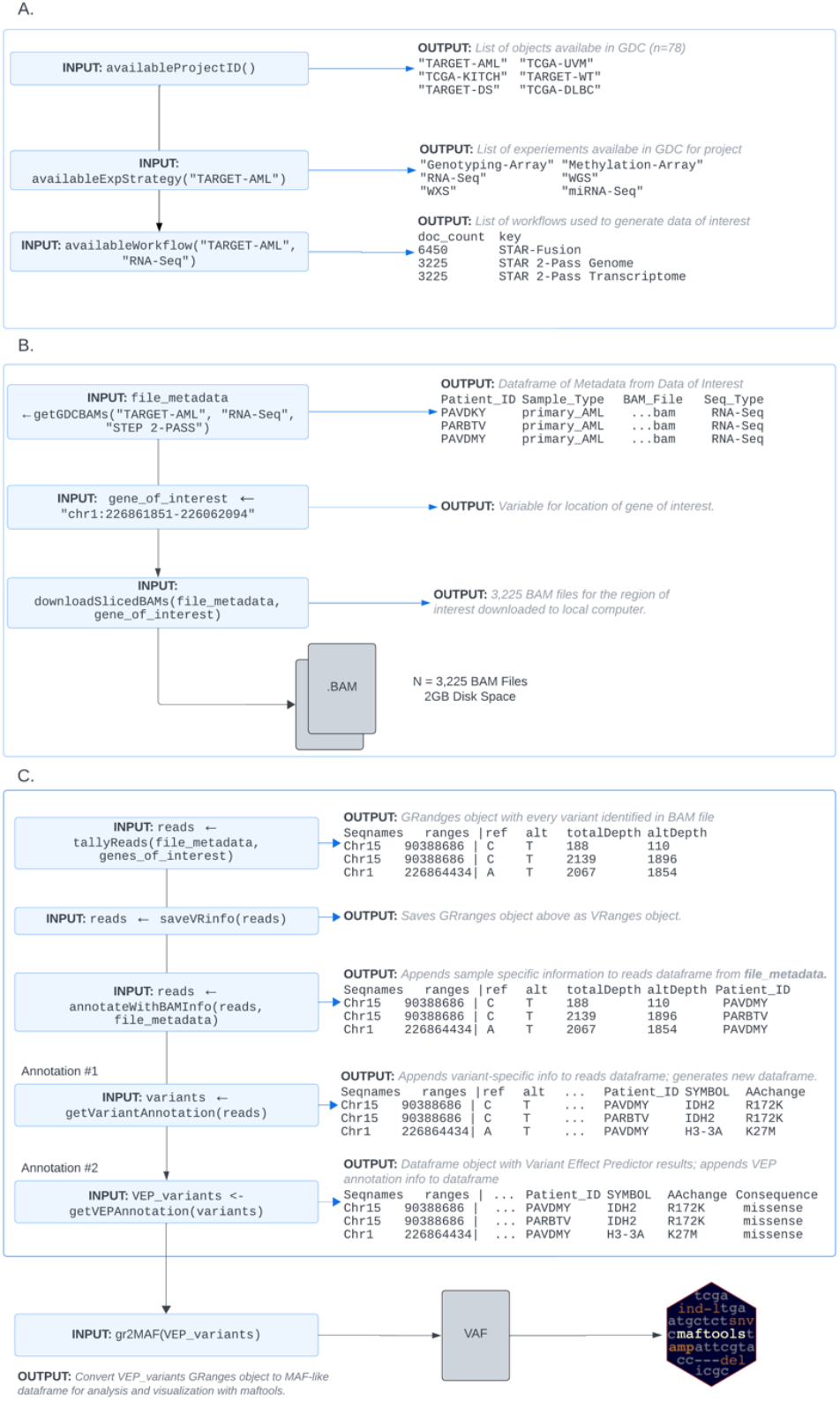
*bamSliceR* Workflow and Functionality for querying, downloading, tallying and annotating variants from BAM files. A) Identify Data Available via GDC Suitable for your Query. B) Download Sliced-BAM files from GDC. C) Variants Tallying & Annotation.

### 2.1 Query Availabile Projects & Download Data of Interest

***bamSliceR*** integrates the GDC API within the R statistical programming environment to enable users to query databases (Figure 2A), remotely BAM-slice genomic regions of interest, and automatically import those sequences for local variant analysis (Figure 2B). We have simplified the querying process into a single function, getGDCBAMs(), to collect metadata of BAM files from the project of interest. Three pieces of information, including the project ID, experimental strategy (ex. “RNA-Seq”, “WGS”), and alignment workflow, are required to locate the BAM files on the GDC portal, which can be obtained using the availableProjects(), availableExpStrategy(), and availableWorkflow(), functions respectively. This workflow allows users to easily select relevant cohorts of patients, sample types, and sequencing data types(WGS/WXS/RNA) according to their specific research question. Based on the criteria specified, getGDCBAMs()will generate a data frame containing metadata for matched BAM files, including patient IDs, sample type(s), sequence type (RNA or DNA), and any other information related to the BAM files of interest. Additionally, the BAM Slicing functionality provided by GDC API requires the user to specify a genomic region(s) of interest with syntax like “chr1:226861851-226062094”. If a user is unfamiliar with the specific coordinates of their gene of interest, bamSliceR provides users with built-in functionality to create a list of exon regions as a vector of strings with the getGenesCoordinates() function. After identifying gene coordinates of interest, bamSliceR users can use the downloadSlicedBAMs() function to download sliced BAM files of the specific genomic regions of the selected cohort.

### 2.2 Tally Variants from BAMs

***bamSliceR*** uses gmapR (Barr C., *et al*. 2016) and VariantTools (Lawrence & Gentleman 2017) to tally read counts of variant alleles and VAF is calculated by altDepth/totalDepth (Figure 2C). A *gmapGenome* object must be created from either a FASTA file or *BSgenome* object that represents the indexed reference genome for use with the GMAP suite of tools. For genomic-aligned BAM files, we can use the *BSgenome* object containing full genomic sequences for humans provided by UCSC as Bioconductor packages, such as *BSgenome*.*Hsapiens*.*UCSC*.*hg38*, which is based on assembly GRCh38.p14. For the transcriptome-aligned BAM files, we recommend using the transcriptome FASTA file from the GENCODE project, which provides well-curated reference files. Tallying reads is the most computationally intense step, especially for memory usage. Similar to BAM slicing, users can specify the regions as a *GRanges* object to tally the reads to speed up the process. Moreover, tallyReads() supports parallel computing across genomic ranges and BAM files, provided a machine with multiple nodes is available. This function includes two arguments (*BPPARAM, parallelOnRangesBPPARAM*) that allow users to configure the number of nodes for parallel processing on either genomic ranges or BAM files, offering flexible control over the efficiency depending on the number of genomic ranges and BAM files. For example, to tally reads on thousands of BAMs but only a few gene regions, to maximize the efficiency, a user can specify more nodes on parallelizing computing on BAM files and fewer nodes on *GRanges* regions.

For the single nucleotide variant (SNV), the readDepth is straightforward: it represents the number of reads covering that single nucleotide position, which can be accurately tallied using tallyReads(). However, the small insertion-deletion variant (INDEL) that spans multiple bases is ambiguous when estimating the reference allele depth. To support INDEL VAF calculation, fixIndelRefCounts() will rescan the BAM files at the regions of INDELs to estimate the depth of reference allele. Users can choose to calculate the depth at the start of the INDEL or calculate the mean coverage of regions that flank the INDEL. ***bamSliceR*** uses *Rsamtools* pileup()to pile up the read counts of each allele within the given ranges and given the same min_mapq and min_base_quality, pileup() and tallyReads() should yield the same read counts for a variant.

### 2.3 Variant Annotation & Visulization

A *VRanges* object will be generated from tallying reads from BAM files containing all the putative variants. AnnotateBAMInfo() will then match the metadata of BAM files back to the corresponding variants. Additionally, getVariantAnnotation() provides the functionality to predict the amino acid changes and consequences of variants using the Bioconductor package *R/VariantAnnotation* (Obenchain, *et al*. 2014) (Figure 2C). Similar to tallying reads, sequence resources either as *BSgenome* objects (for genomic-aligned BAM files) or FASTA files (for transcriptome-aligned BAM files) must be provided by the user. Additionally, the *TxDb* object that serves as the annotation database is required for amino acid change prediction of the variants. UCSC provides annotation databases from different species as Bioconductor packages, such as *TxDb*.*Hsapiens*.*UCSC*.*hg38*.*knownGene* for humans is ready for annotating variants with genomic coordinates.

UsingVariant Effect Predictor (VEP), a user can curator detailed annotations of the effects of variants on transcripts and genes (McLaren, *et al*.*2*016). Notably, VEP predicts the functional impact of variants, identifying whether they cause missense, nonsense, synonymous, or frameshift mutations, and their potential effects on protein function based on scores from various tools (e.g., SIFT, PolyPhen). Using gr2vrforVEP(), ***bamSliceR*** allows users generate an input VCF file containing candidate variants from patients for use with VEP. After running VEP locally, users can parse the consequence (CSQ) column in the resulting VCF file to annotate the variants with information provided by VEP with the getCSQFfromVEP() function (Figure 2C).

To enable downstream analysis and visualization of these annotated results, ***bamSliceR*** allows users to export the annotated variant data into a Mutation Annotation Format (MAF) file, specifically designed for compatibility with the Bioconductor package *R/ maftool*s (Mayakonda, *et al*. 2018), which is widely used for the analysis and visualization of somatic mutations in cancer genomics studies (Figure 2C).

The exported MAF file includes essential information such as gene names, variant classifications, amino acid changes, and functional impacts, making it ready for further analysis (Figure 2C). By converting variant data into MAF format, researchers can seamlessly integrate with *R/maftools* to generate visualizations such as oncoplots, lollipop plots, and mutational signature analyses, which are crucial for interpreting the mutational landscape in cancer studies (Mayakonda, *et al*. 2018). This feature simplifies data preparation, letting users focus on analysis rather than formatting.

### 2.4 Transcript-Aware Annotation

***bamSliceR*** facilitates transcript-aware variant annotation from transcriptome-aligned BAM files. The workflow is similar to methods described above with genomic-aligned BAM files, but incorporates additional utilities, including: generation of transcript annotation files compatible with Bioconductor’s built-in toolset and finding the coordinates of all transcripts corresponding to the genomic coordinates of a variant (Figure 1B).

### 2.2 Mapping Transcripts Coordinates of Features in GFF3 File

General Feature Format version 3 (GFF3) is a standard format used to describe genomic features like genes, transcripts, exons, and coding sequences (CDS) (Figure 3). In a GFF3 file, the features of genes are organized in a hierarchical structure that reflects their biological relationships (Figure 3). Bioconductor packages like *rtracklayer* and *GenomicFeatures* can read GFF3 files to import genomic features into R as *GRanges* objects. Moreover, *GenomicFeatures* can use GFF3 files to create the previously mentioned *TxDb* object for variant annotation.

**Figure 3.**
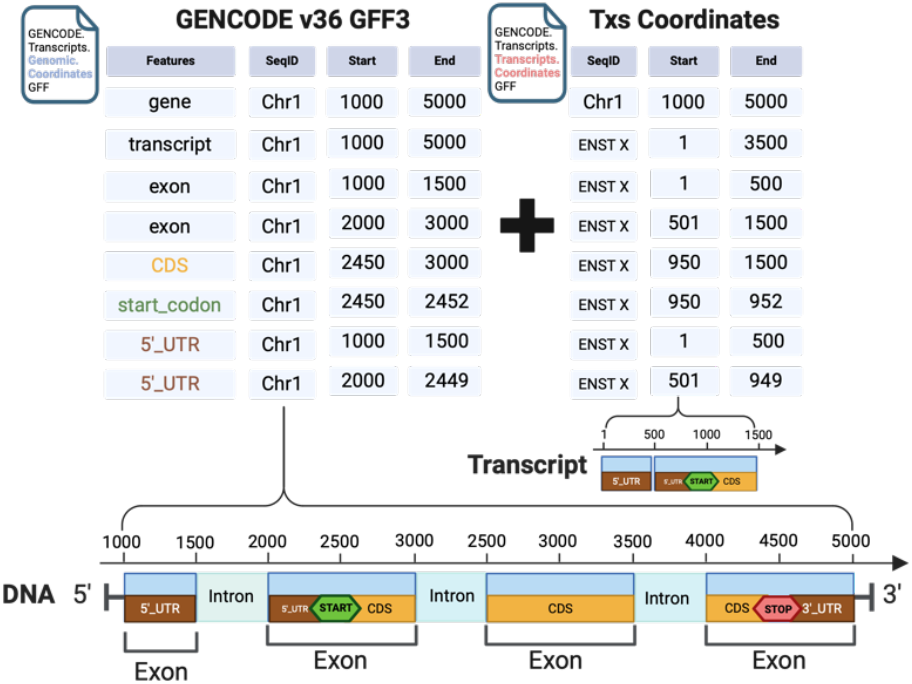
Scheme to Calculate Transcript Coordinates of Genomic Features in GFF3 file.

However, conventional GFF3 files from GENCODE contain gene annotations on genomic coordinates incompatible with the Transcriptome-aligned BAMs, which lack the genomic coordinates when mapping reads to reference transcript sequences. The *VRanges* object generated from tallying reads of Transcriptome-aligned BAM files records transcripts coordinates with unique transcript IDs as sequence names provided by the Ensembl database. To annotate the tallied variants from a transcriptome BAM, ***bamSliceR*** can parse each genomic feature type of a gene in GFF3 and then calculate the coordinates relative to transcripts for each feature entity based on the given genomic coordinates (Figure 3). The newly-generated GFF3 file preserves the hierarchical structure of features and includes coordinate information for both genomic and transcript for each feature entity. Downstream analysis of variants in the context of the transcriptome relies on the modified GFF3 file, such as variant annotation and matching results between genomic- and transcriptome-aligned BAMs.

### 2.3 Finding Equivalence Class of Transcripts for Variants

When a genetic variant, such as an SNP or an INDEL, occurs at a specific genomic position, it may impact multiple transcripts that overlap at that position. The set of transcripts that share the exact genomic coordinates for the given variant can be defined as an equivalence class of the variant. ***bamSliceR*** incorporates an additional step to identify and re-tally read counts for the equivalence class of each variant before performing transcript-aware annotation. This step is essential for accurately annotating transcripts overlapping with a variant but with few or no mapped reads, which might otherwise be omitted or filtered out. Using the modified GFF3 file mentioned above, ***bamSliceR*** provides the function getDisjoinOverlapBins() to generate disjointed bins for each gene, containing both transcriptome and genomic coordinates of overlapping transcripts. The bins that overlap with a given variant are identified, and the equivalence class for the variant is determined by finding the intersection of all overlapping bins. The variant’s coordinates relative to each transcript in the equivalence class that were previously missing are then calculated. Then, ***bamSlicer*** provides the function fixMissingTxs()to re-scan corresponding BAM files to tally the read counts at these loci (Figure 4).

**Figure 4.**
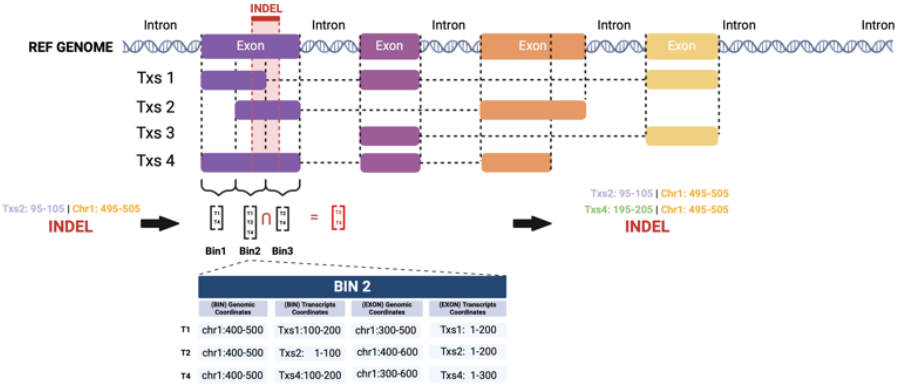
Schematic for Determining Equivalence Classes of Transcripts for INDEL variants. To identify equivalence classes of transcripts for INDEL variants, the process beings by disjoining exon regions for each gene in the GFF3 file based on their genomic coordinates. For instance, a single exon may be split into multiple disjoint bins. In the provided example, the first exon generates three bins: Bin1 [T1, T2], Bin2 [T1, T2, T3], and Bin3 [T2, T3]. For each bin, the genomic and transcript-level coordinates of both the bin and the corresponding exon are tabulated (e.g., see the detailed example for Bin2). When analyzing an INDEL, all bins overlapping with the variant’s genomic coordinates are identified. Finally, the transcripts containing the INDEL are determined by intersecting the sets of transcripts associated with the overlapping bins.

### 2.4 Annotation of Variants in the Context of the Transcriptome

After tallying read counts across all possible transcripts for each variant, ***bamSliceR*** facilitates variant annotation. Similar to annotating variants from genomic BAM files, getVariantAnnotation.Txs() uses the *VariantAnnotation* package to predict amino acid changes and variant consequences. This process relies on annotation databases and sequence resources compatible with transcriptome-aligned BAMs. Specifically, it requires a modified GFF3 file containing genomic and transcriptomic coordinates for each gene’s transcript features to generate the *TxDb* object and a FASTA file with nucleotide sequences of the transcripts. The getGenCodeAnnotation.Txs() can map variant loci to the features described in the GENCODE GFF3 annotation file. This process covers not only coding sequence (CDS) regions but also focuses on UTRs, START codons, and STOP codons. Additionally, ***bamSliceR*** aligns the transcriptomic coordinates of each variant with their corresponding genomic coordinates. In practice, ***bamSliceR*** provides a wrapper function, *getVariantAnnotationForTxs()*, that can retrieve annotation results from both sources, ensuring comprehensive transcript-aware variant annotation.

### 2.5 Genomic Data Commons Controlled Access

Access to the GDC is not limited solely to registered users; rather, it accommodates a broad spectrum of user-types engaged in cancer research and related endeavors. Those seeking access to GDC data must complete a straightforward registration process, facilitated through the official GDC portal ([Getting Started - GDC Docs](https://docs.gdc.cancer.gov/Data_Portal/Users_Guide/Getting_Started/) (cancer.gov)). In order to maintain security of genomic data, the GDC requires users to authenticate their credentials using a unique GDC token. To obtain a GDC token, registered users must navigate to the GDC portal and follow the prescribed guidelines for token generation and authentication. Once the unique GDC token has been generated, users are prompted to save the downloaded GDC token in a designated file within their user home directory, denoted as “.gdc_token”, which is used by downloadSlicedBAMs()to allow for access to GDC files.

## 3 Results

### 3.1 Reduction of Disk Space Required for Analysis by *bamSliceR*

The 3,225 RNAseq BAM files obtained from 2,281 subjects in TARGET-AML are approximately 39TB. Using ***bamSliceR*** to slice the reads covering exons of 11 Histone 3 genes (*H3F3A, HIST1H3A, HIST1H3H, HIST1H3I, HIST1H3J, HIST1H3B, HIST1H3C, HIST1H3D, HIST1H3E, HIST1H3F, HIST1H3G*) and genes encoding epigenetic factors *(IDH1/2, DNMT3A, RUNX1, ASXL1/2, TET1/2*) that collectively span 152,026 basepairs of non-contiguous genomic space reduced the total size of BAM files to 62GB. Furthermore, slicing the reads covering a single human *MLLT1* (ENL) gene span of 73,594 base pairs only requires 2GB of total disk space for 3,225 BAM files (Figure 5). This drastic reduction of disk space necessary to perform variant analysis makes local analysis of data from large cohorts much more accessible to a traditional researcher.

**Figure 5.**
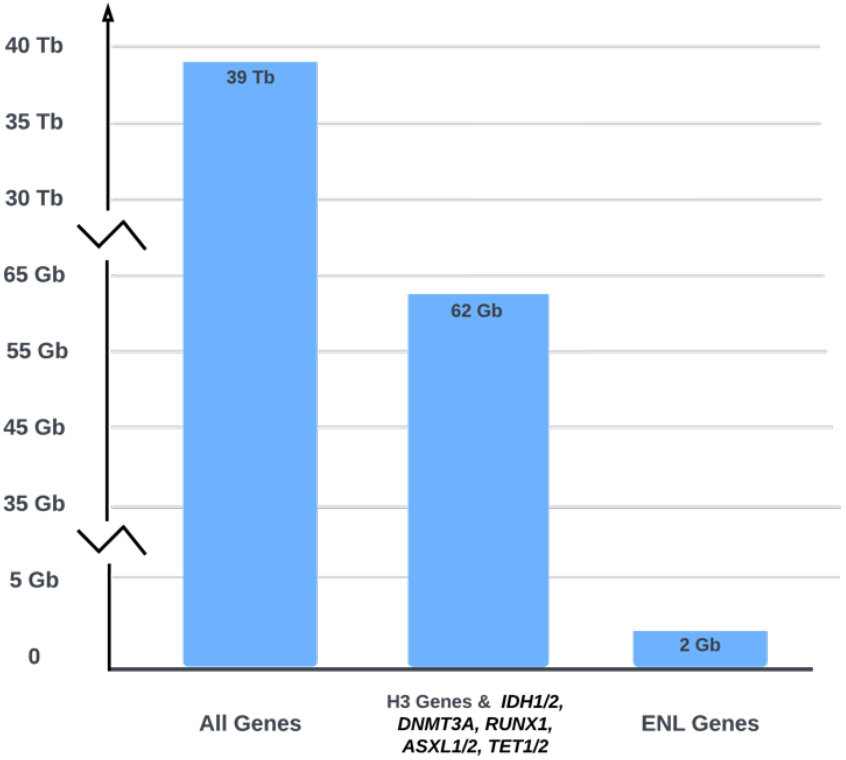
bamSliceR is designed to significantly reduce the space required for analyzing BAM files by slicing them at specified genomic regions

### 3.2 Genomic Data Alignment Case Study: Oncohistone Variants in Pediatric Leukemia

Pediatric acute myeloid leukemia (AML) is a genetically heterogeneous and often lethal disease (Li, *et al*. 2016). Most cases reveal few, if any actionable sequence variants, with a remarkably low mutation burden and a preponderance of diverse structural variants (Bolouri, *et al*. 2018). Nevertheless, molecular features can drive treatment decisions. For example, most patients with *DEK::NUP214* fusions harbor co-occurring *FLT3* internal tandem duplications. Targeting this aberration revealed the immunogenicity of the fusion, and when combined with stem cell transplantation, has improved 5-year survival from <10% to >85% in young patients (Tarlock, *et al*. 2021). Unfortunately, identifying actionable sequence variants from WGS is challenging both technically and statistically, as the population of AML patients is small relative to common diseases and the total number of pediatric cases with WGS is in the low hundreds. The analysis of expressed sequence variants from RNA-seq reads is therefore quite attractive.

Mutations of histone H3.3 lysine 27 (H3K27) to methionine (M, H3K27M) were originally documented in high-grade midline glioma, where they frequently accompany *TP53* mutations (Schwartzentruber, *et al*. 2012). More recently, mutations of H3.1 K27 to isoleucine (I) or methionine (H3K27I/M) were documented in adult AML patients (Lehnertz, *et al*. 2017) and in pre-leukemic stem cells from patients who went on to develop secondary AML(Boileau, *et al*. 2019). However, the genetic context, age groups (histone mutations have not previously been reported in pediatric AML patients), mechanism, and clinical impact of H3K27 mutations in myeloid leukemogenesis remains poorly understood at best.

To investigate the genetic landscape of H3K27M in AML across age groups, we examined 11 histone 3 genes and genes encoding epigenetic factors that are frequently mutated in AML patients (*IDH1/2, DNMT3A, RUNX1, ASXL1/2*, and *TET1/2*). These genes collectively span 152,026 bp of non-contiguous genomic space. Using the *bamSliceR* pipeline, we automatically processed 3,225 and 735 RNA-seq BAM files obtained from 2,281 and 653 subjects in the TARGET-AML (pediatric) and BEAT-AML (adult) cohorts, respectively. This step alone reduced the total size of the BAM files of the TARGET-AML cohort from 39TB to 62GB (Figure 5), while still retaining all the essential genetic Information required to perform an epidemiological study of these rare AML mutations. We identified 9 pAML and 7 adult AML patients that harbored a K27M mutation based on Variant Allele Frequency (VAF >0.15), total read depth (>8), and WGS data where available (Supplementary Table S1). We found H3K27M mutations on both replication-coupled H3.1 and replication-independent H3.3, with most mutations occurring in the H3.1 gene (Supplementary Table S1). The incidence of H3K27M in pediatric AML is lower (∼0.3 %) than in adult AML (∼0.8%; Supplementary Table S2), and we confirmed the existence of H3K27 variants in normal karyotype induction failure patients (Table 1).

**Table 1.**
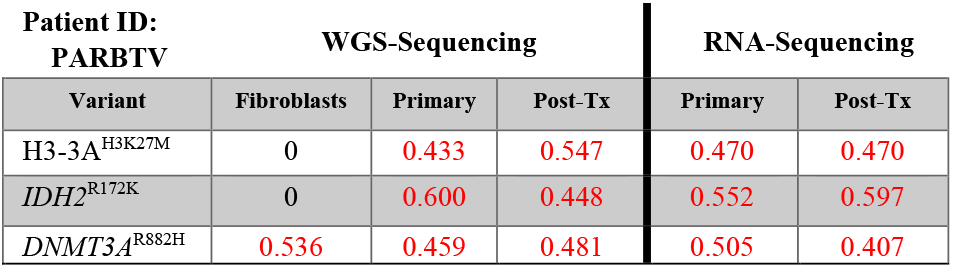
Clonal mutations observed in TARGET AML subject *PARBTV* from WGS and RNAseq Data. Variant allele frequency of mutations H3F3A K27M, IDH2 R172K (somatic), and DNMT3A R882H (germline) of a pediatric patient from the TARGET-AML cohort with both RNA-seq and WGS data from two timepoints (diagnosis and post-treatment).

We then used ***bamSliceR*** to generate VAF distribution plots for each mutation and estimate their clonal status. For example, we see that *IDH2* ^R172K^ and *DNMT3A*^R132H^ mutations are persistent clonal events (VAF ∼50%) in the pAML cohort, mutations that were previously believed to only occur in adult AML patients (Supplementary Fig S1C; S2). We discerned that the K27M mutation in *H3C2, H3C3, H3C4, H3C11* (H3.1) and *H3F3A* (H3.3) genes are always clonal (Supplementary Fig S3), consistent with their occurrence in pediatric high-grade glioma. Oncoplots and mutual exclusivity analysis showed that *H3F3A* K27M (H3.3) and *IDH2* mutations co-occur (p<0.05) (Supplementary Fig S1D). Interestingly, two pAML patients harboring *H3F3A* K27M and *IDH2*^R172K^ mutations failed induction therapy, suggesting that the two mutations may synergize to cause chemoresistance. For pAML patients where RNA-seq and WGS data were both available, we used ***bamSliceR*** to confirm that the somatic *H3F3A* K27M and *IDH2*^R172K^ mutations are constantly expressed at high levels and with low allelic bias, consistent with DNA sequencing results (Supplementary Table S1; Fig S1D). To facilitate in-depth studies of patients with samples at multiple disease stages, ***bamSliceR*** includes functionality to identify and subset the matched subjects and data files. Using this function, we found that one pAML patient with somatic *H3F3A* K27M and *IDH2*^R172K^ mutations also harbored a germline *DNMT3A*^R882C^ mutation throughout disease progression (Table 1).

Taken together, these data suggest that H3K27M is a clonal mutation that may synergize with metabolic and epigenetic variants (e.g. *IDH2*^R172K^ and *DNMT3A*^R882C^) to drive aggressive and refractory pAML. *IDH* mutations have been thought mutually exclusive with H3K27 variants. Not only do they co-occur, *IDH2* mutations are in fact enriched for H3K27 mutant cases. By using RNA-seq data to expand our sample size and automating the alignment and variant calling process in bamSliceR, we document statistically and clinically significant co-occurrence of oncometabolic *IDH2* variants with high-risk H3K27 pAML mutant cases, yielding testable hypotheses and new translational avenues (Thomas, *et al*. 2023) for a subset of AML patients at high risk of treatment failure.

### 3.3 MLLT1 YEATS Domain Indels in Pediatric Tumors

The ENL protein (encoded by the *MLLT1* gene) is a subunit of the super elongation complex (SEC) involved in transcriptional elongation during early development. Small, in-frame INDEL mutations in the YEATS domain of *MLLT1* were first identified in Wilms tumors (Perlman, *et al*. 2015). Further work revealed that indels in the YEATS domain alter chromatin states, dysregulating cell-fate control and driving tumorigenesis (Wan, *et al*. 2020). These same indels can transform hematopoietic cells by mitigating polycomb silencing, but only one case of pediatric AML harboring this type of mutation has previously been documented (http://cancer.sanger.ac.uk/cosmic).

As a second case study for the efficacy of our package, we used ***bamSliceR*** to analyze the TARGET RNA-seq and WGS data for any evidence of ENL (*MLLT1*) YEATS domain mutations. We discovered three pAML patients carrying *ENL*-YEATS mutations, with one pAML patient exhibiting allelic bias similar to that observed in Wilms tumor patients (Table 2). We also identified putative indel mutations near the YEATS domain that have not been detected in Wilms tumor patients (Supplementary Table S3). These new results lead to the biologically testable hypothesis that *ENL*-YEATS mutations primarily upregulate *HOXA* gene expression in pAML, similar to their role and function in favorable histology Wilms tumors (Gadd, S. *et al*. 2017).

**Table 2.**
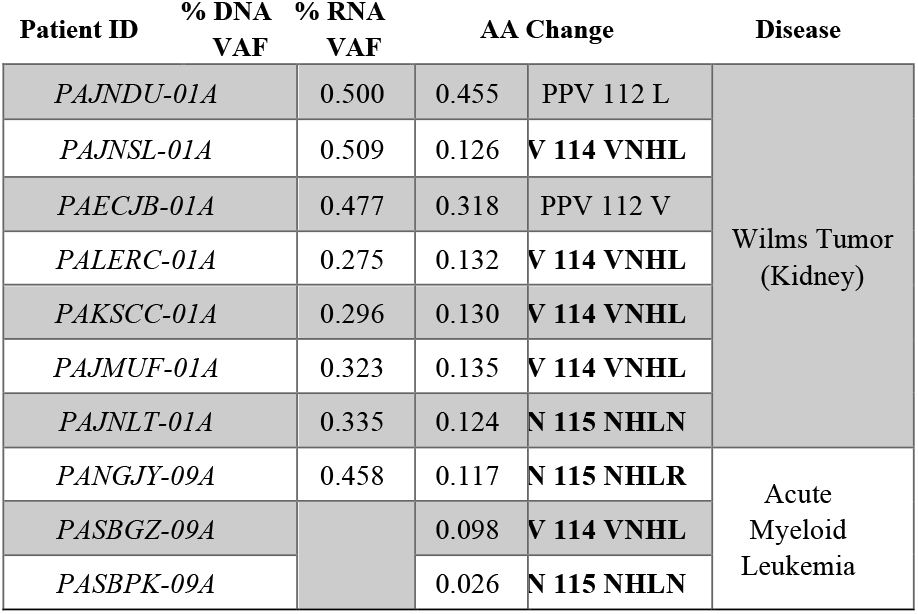
Rare ENL YEATS Domain Mutations in pAML patients. Allelic bias based on Variant Allele Frequency of in-frame insertion mutations in ENL from Wilms tumor and AML patients where both WTS and WGS data were available.

### 3.4 Re-Diagnosing Complex Structural Variants: Detection of UBTF-ITD in Previously Annotated Samples

The primary functions of *bamSliceR*—including variant pileup and annotation— are fully compatible with locally stored BAM files. We demonstrate the application of *bamSliceR* on publicly available RNA-seq data from the Leucegene cohort, which includes 452 transcriptomes of AML patients with an average depth of 204 million reads per sample. This analysis specifically focuses on larger structural variants detection in AML patients. We highlight new diagnostic results of a known internal tandem duplication (ITD) event on exon

13 of upstream binding transcription factor (UBTF) that has recently been characterized as a novel prognostic genetic variation in AML, particularly in relapse cases (Umeda, *et al*. 2022). Umeda, M. *et al* (2022) reported 9 pediatric AML patients with UBTF-ITD in the TARGET dataset, identified using CICERO (v1.7.0) (Tian, *et* al. 2020) and a novel soft-clipping ratio method (Umeda, *et al*. 2022). These cases are accessible via GDC portal and can serve as positive-control when deteceing UBTF-ITD events (Supplementary Figure S4). Using the BAM-Slicing pipeline utilized samtools to extract reads at given genomic ranges, we selectively retain reads aligned to UBTF locus, reducing the overall data size to ∼1G BAM files from an initial dataset of 4.17 TB, optimizing downstream analysis on high-depth RNA-seq data.

The UBTF-ITD typically involves in-frame insertions, ranging from 45 to 95 base pairs, at the 3’ end of exon 13. These insertions lead to clusters of soft-clipped nucleotide sequences near this region (Umeda, *et al*. 2022). Utilizing bamSliceR to tally the soft-clipped read counts at each locus, we identified two patients in the Leucegene cohorts with multiple instances of high soft-clipped read counts around the UBTF-ITD hotspots (Figure 6). We then performed manual examination of the RNA-Seq BAM files of these patients to validate our method’s efficacy in detecting true-positive of UBTF-ITD events (Figure 6). The initial discovery of UBTF-ITD in 2022 underscored the challenges in detecting these rare and complex mutations. However, expanding the sample size by screening for known mutations in available cohorts should be feasible and efficient. Here, we demonstrate the utility of *bamSliceR* to transform publicly available sequencing data into actionable diagnostic insight, even for complex structural variants.

**Figure 6.**
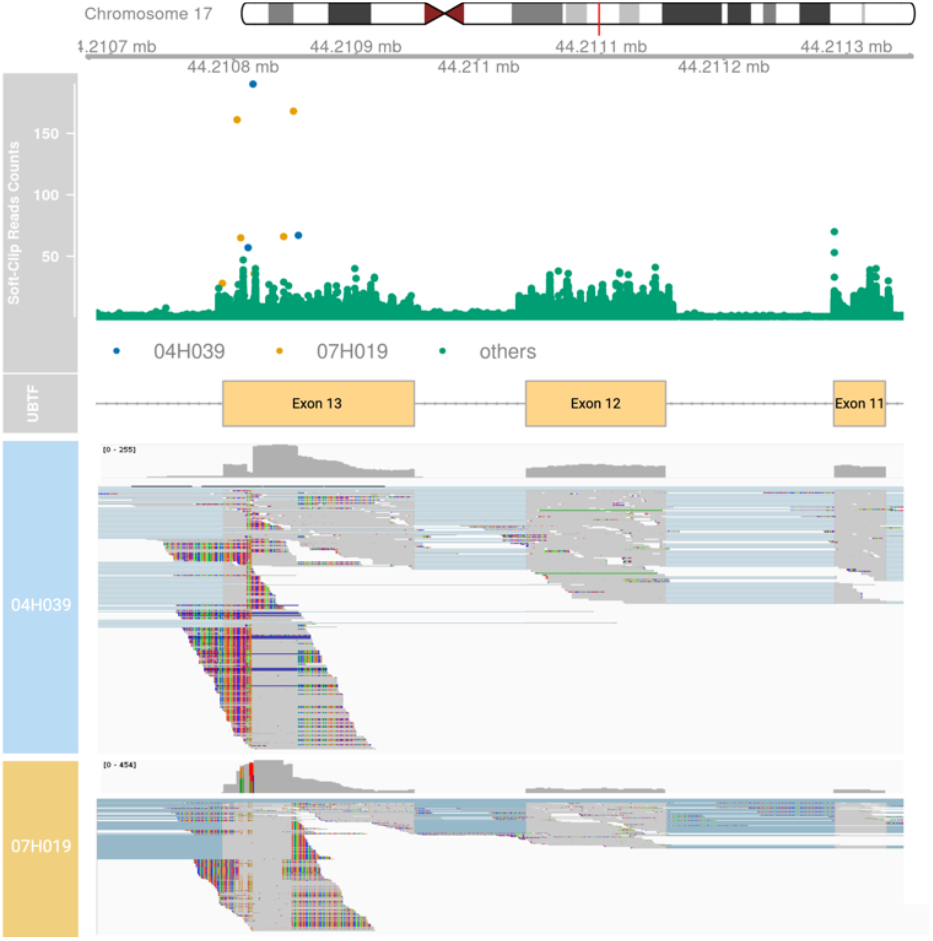
Detection of UBTF-ITD in the Leucegene Project. Top) Plot of soft-clip read counts (sc-counts) within chr17:44210679-44211356 (GRCh38) of 452 patients in the Leucegene Project. Two patients (04H039 and 07H019) show high soft-clipped read counts at hotspots of UBTF-ITD. 04H039 has 57, 190, 67 sc-counts at chr17:44210812, chr17:44210816 and chr17:44210853 respectively (Highlight in Blue dots). 07H019 has 161, 65, 66, 168 sc-counts at chr17:44210803, chr17:44210806, chr17:44210841, and chr17:44210849, respectively (Highlight in Yellow dots). Manual examination of RNA-seq BAM files of 18 patients with a maximum sc-counts > 25 within the ranges show no UBTF-ITD. Bottom) Integrative Genomics Viewer (IGV) visualization showing UBTF-ITD with soft-clipped reads and increased coverage in UBTF exon 13 in patients 04H039 and 07H019.

## 4 Discussion

We developed ***bamSliceR*** to address two practical challenges: resource-sparing identification of candidate subjects, and variant detection from aligned sequence reads across thousands of controlled-access subjects. The GDC BAM Slicing API (https://docs.gdc.cancer.gov/API/Users_Guide/BAM_Slicing/) is a practical and well-documented REST API for this purpose. Yet, we find relatively little published work that uses the GDC API, even in studies where specific candidate variants are evaluated. Instead, reliance upon variant calls from existing studies, or transfer and recall of variants from raw sequence data, is often documented. The former assumes a single best method for variant detection fits all experimental designs (an assumption contradicted by many benchmarks[*cite*]). The latter is grossly inefficient.

In clinical genetic analysis, direct evaluation of fragment-level evidence for a candidate variant is routine, regardless of the confidence level a variant calling pipeline may assign to a putative genetic variant. The same raw material is available via controlled access across many population-scale projects representing billions of dollars in public funding. The relatively simple toolkit we provide here extends this practice in an efficient and user-friendly way to the Bioconductor ecosystem, significantly expanding the pool of potential subjects for rare variant detection in rare diseases. Importantly, the same well-documented API implemented by the GDC is feasible for Common Fund datasets, such as the Gabriella Miller Kids First! (GMKF) and INCLUDE projects, and can be further extended to transcriptome-indexed reads and open access resources like SRA.

Support for efficient retrieval of range-based queries is a key feature of the SAM/BAM format and htslib/htsget implementations. Authentication and authorization create challenges for straightforward usage in controlled access data, but as the GDC API and its downstream users illustrate, this challenge can be overcome, and is increasingly important as more WTS and WGS data from population-scale biobanks emerge. We present ***bamSliceR*** as a concrete example of what can be accomplished by wedding clinical and genomic data management processes to an efficient, standardized API. Our hope is that it will enable users to perform previously challenging evaluation of raw data evidence for genetic and genomic variants at scale, and that user uptake will spur further expansion of support for coordinate-based queries of sequence databases.

## Acknowledgements

Computation for the work described in this paper was supported by the High Performance Cluster and Cloud Computing (HPC3) Resource at the Van Andel Research Institute. We thank Darrell Chandler for his expertise and insight throughout all aspects of the study, and for his assistance in copy-editing the manuscript. Some of the results obtained in this publication are based upon data generated by the Leucegene group primarily located at IRIC in Montreal, Canada and supported by Genome Canada and Genome Québec. This data was made possible through human AML specimens provided by the BCLQ, Montreal, Canada. To cite a tool developed by the Leucegene Project’s team, please follow the instructions provided by each specific tool.

## Data availability

The data underlying this article are available from the NCBI Sequence Read Archive (SRA) (Leucegene project SRP056295, GEO GSE67040) and the database of Genotypes and Phenotypes (dbGaP) projects phs000218 (NCI/COG TARGET) & phs001657 (BEAT-AML), which can be accessed directly via the NCI Genomic Data Commons (GDC) or via bamSliceR with the appropriate data use authorization (granted via dbGaP and ERA commons).

## Supplementary data

Supplementary data are available at *Bioinformatics Advances* online.

