## Supplemental materials and figures (v3) for "*bamSliceR*: a Bioconductor package for rapid, cross-cohort variant and allelic bias analysis"

**Supplemental Data**

### Table S1. Co-occurrence status of *H3K27M* and *IDH2* mutations in TARGET-AML and BEAT-AML.

Identification of 9 pAML patients that harbored K27M mutation on Histone 3 genes based on evidence of VAF > 0.15 and total read depth > 8 and whether the mutation can be captured by both RNA-seq and WGS data. Co-occurrence of *H3F3A* (H3.3) *K27M* and *IDH2 R172K* are shown in 2 pAML patients. Co-occurrence of *H3K27M* (H3.3 & H3.1) and *IDH2 R140Q/R172K* are shown in 5 adult AML patients.

|  |  | **H3K27M** | | | | | **IDH2** | | | | |
| --- | --- | --- | --- | --- | --- | --- | --- | --- | --- | --- | --- |
|  | **CASE_ID** | **SYMBOL** | **DNA-vaf-T1** | **DNA-vaf-T2** | **RNA-vaf-T1** | **RNA-vaf-T2** | **SYMBOL** | **DNA-vaf-T1** | **DNA-vaf-T2** | **RNA-vaf-T1** | **RNA-vaf-T2** |
| **H3.3** | PARBTV | H3F3A | 47% | 48% | 47%% | 47% | R172K | 38% | 48% | 56% | 59%% |
|  | PAVDMY | H3F3A |  |  | 49% | 47%/47% | R172K |  |  | 39% | 40%/48% |
| **H3.1** | PAPVGE | HIST1H3C |  |  | 58%% |  | - |  |  | - |  |
|  | PAUZTH | HIST1H3J^K27I^ |  |  | 0% | 38%% | - |  |  | - | - |
|  | PAUUPR | HIST1H3I |  |  | 0% | 35%% | - |  |  | - | - |
|  | PAKWCU | HIST1H3D |  |  | 17%% |  | - |  |  | - |  |
|  | PAXFAG | HIST1H3C |  |  | 54%% |  | - |  |  | - |  |
|  | PAXKAL | HIST1H3I |  |  | 50% |  | - |  |  | - |  |
|  | PATFGK | HIST1H3C |  |  | 50% |  | - |  |  | - |  |

|  |  | **H3K27M** | | | | **IDH2** | | | |
| --- | --- | --- | --- | --- | --- | --- | --- | --- | --- |
|  | **CASE_ID** | **SYMBOL** | **DNA-vaf** | **RNA-vaf** | **TS-vaf** | **SYMBOL** | **DNA-vaf** | **RNA-vaf** | **TS-vaf** |
| **H3.3** | 2148 | H3F3A | 29% | 47% |  | R140Q | 40% | 52% |  |
| **H3.1** | 2354 | HIST1H3C | 34%/33% | low-exp |  | R172K | 36%/31% | 47% |  |
|  | 2429 | HIST1H3B | 33%/47% | 35.29%/100% |  | - | - | - |  |
|  | 2498 | HIST1H3C | 37% | 38% |  | R140Q | 48% | 47% |  |
|  | 2530 | HIST1H3B | 40% | low-exp |  | R172K | 46% | 49% |  |
|  | 2611 | HIST1H3D |  | 58% | 39% | - |  | - | - |
|  | 2721 | HIST1H3B |  |  | 36% | R172K |  |  | 36% |

#### **Table S2** *H3K27* variants: published work (n=1049) and TARGET/BEAT-AML cohorts (n = 2934, via *bamSliceR*)

Lehnertz et al. *Blood* *2017* documented 2 adult AML patients. Boileau et al. *Nat Commun 2019* documented 4 adult AML patients. We documented 16 AML patients from TARGET and BEAT AML cohorts.

| **Cohort** | **Cohort Size (n)** | **WGS/WXS (n)** | **RNAseq (n)** | **DNA Methylation (n)** | **H3K27M/I (%/n)** |
| --- | --- | --- | --- | --- | --- |
| **Lehnertz et al. *Blood* (2017)** | | | | | |
| **Leucegene** | 415 |  | 415 |  | 0.48%/2 |
| **Boileau et al. *Nat Commun* (2019)** | | | | | |
| **Toronto** | 312 | 312 |  |  | 0.64%/2 |
| **Lebanon** | 122 | 122 |  |  | 0.8%/1 |
| **TCGA** | 200 | 200 | 200 | 200 | 0.5%/1 |
| **Identification of H3K27M in BEAT-AML and TARGET-AML** | | | | | |
| **TARGET 20/21** | 2045 | 365 | **2281** | 2000 | 0.4%/9 |
| **Beat-AML** | 826 | 798 | **653** |  | 0.8%/7 |

#### Table S3

**Human *MLLT1* YEATS domain insertion/deletion variants identified in the pan-TARGET cohort.**

| **Chromosome** | **POS** | **SYMBOL** | **AAchange** | **REFCODON** | **VARCODON** | **REFAA** | **VARAA** | **alt_count** | **totla_count** | **VAF** | **patient_id** |
| --- | --- | --- | --- | --- | --- | --- | --- | --- | --- | --- | --- |
| **chr19** | 6230649 | MLLT1 | V114VNHL | GTG | GTGAACCACCTG | V | VNHL | 5 | 51 | 0.09804 | PASBGZ |
| **chr19** | 6230642 | MLLT1 | H116HLRP | CAC | CACCTGCGCCCC | H | HLRP | 2 | 460 | 0.00435 | PANGJY |
| **chr19** | 6230645 | MLLT1 | N115NPLR | AAC | AACCCCCTGCGC | N | NPLR | 4 | 470 | 0.00851 | PANGJY |
| **chr19** | 6230645 | MLLT1 | N115NHLR | AAC | AACCACCTGCGC | N | NHLR | 55 | 470 | 0.11702 | PANGJY |
| **chr19** | 6230640 | MLLT1 | HL116L | CACCTG | CTG | HL | L | 2 | 134 | 0.01493 | PALHVV |
| **chr19** | 6230646 | MLLT1 | N115NHLH | AAC | AACCACCTGCAC | N | NHLH | 2 | 76 | 0.02632 | PASBPK |
| **chr19** | 6230582 | MLLT1 | LL135L | CTCCTG | CTG | LL | L | 2 | 175 | 0.01143 | PAUMUZ |
| **chr19** | 6230611 | MLLT1 | NP126P | AACCCC | CCC | NP | P | 2 | 370 | 0.00541 | PAUHGM |
| **chr19** | 6230616 | MLLT1 | TFN123N | ACCTTCAAC | AAC | TFN | N | 2 | 48 | 0.04167 | PAWVPZ |
| **chr19** | 6230620 | MLLT1 | TF123F | ACCTTC | TTC | TF | F | 2 | 690 | 0.00290 | PAVDXR |
| **chr19** | 6230574 | MLLT1 | AG138G | GCCGGC | GGC | AG | G | 2 | 175 | 0.01143 | PABYYR |
| **chr19** | 6230570 | MLLT1 | GG139G | GGCGGG | GGG | GG | G | 2 | 1047 | 0.00191 | PAUWZR |

#### **Figure S1**

**Figure S1. *bamSliceR* Data Visualization Functionality**

**A-D** Example of automatically generation of *Oncoplot, survival analysis, VAF distribution* and *Mutual Exclusivity analysis.* **B.** VAF plotting of different mutations of *IDH2* at either *R140* or *R172* ordering by median of VAF. *R140Q* and *R172K* are most prevalent and *R172K* mutation are always clonal which are usually have mean allele frequency around ~50% assuming pure sample (VAF plotting of *H3K27M* and *DNMT3A* are in Supplementary Figure S1 and S2 ). **C.** Kaplan meier curve by grouping samples based on mutation status (WT vs. DNMT3A) in both TARGET and BEAT AML cohorts (n = 2934). **D.** Mutual exclusivity plot by pair-wise Fisher’s Exact test detected *H3-3A K27M* and *IDH2 R172K* are co-occuring mutations (p < 0.0032).

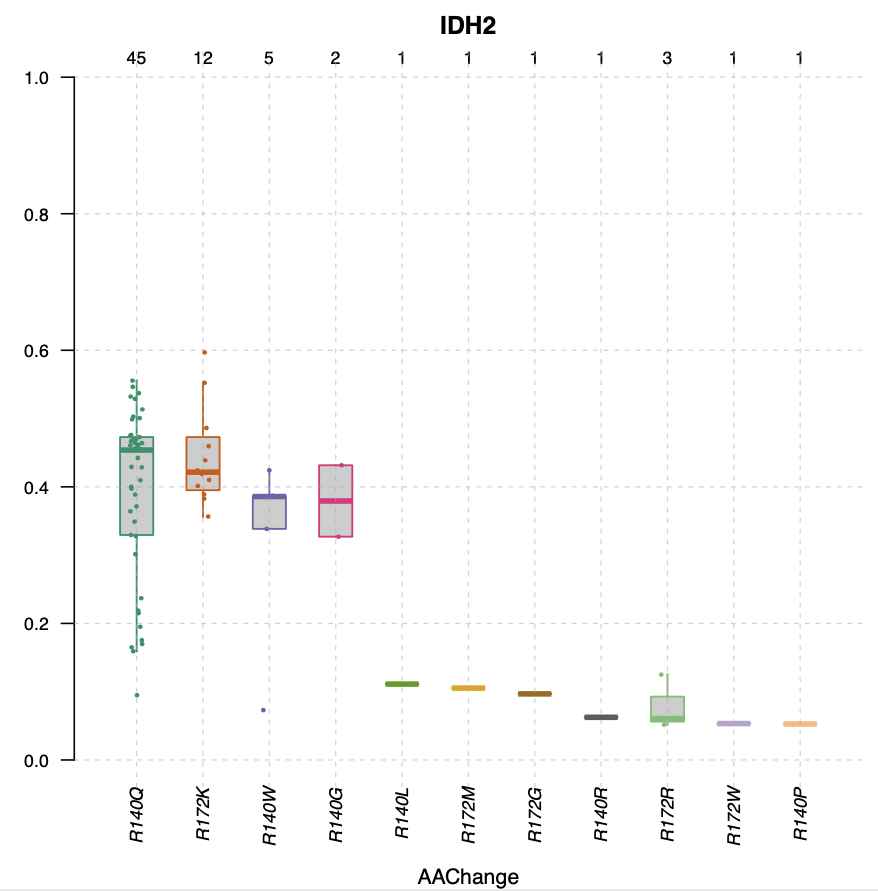

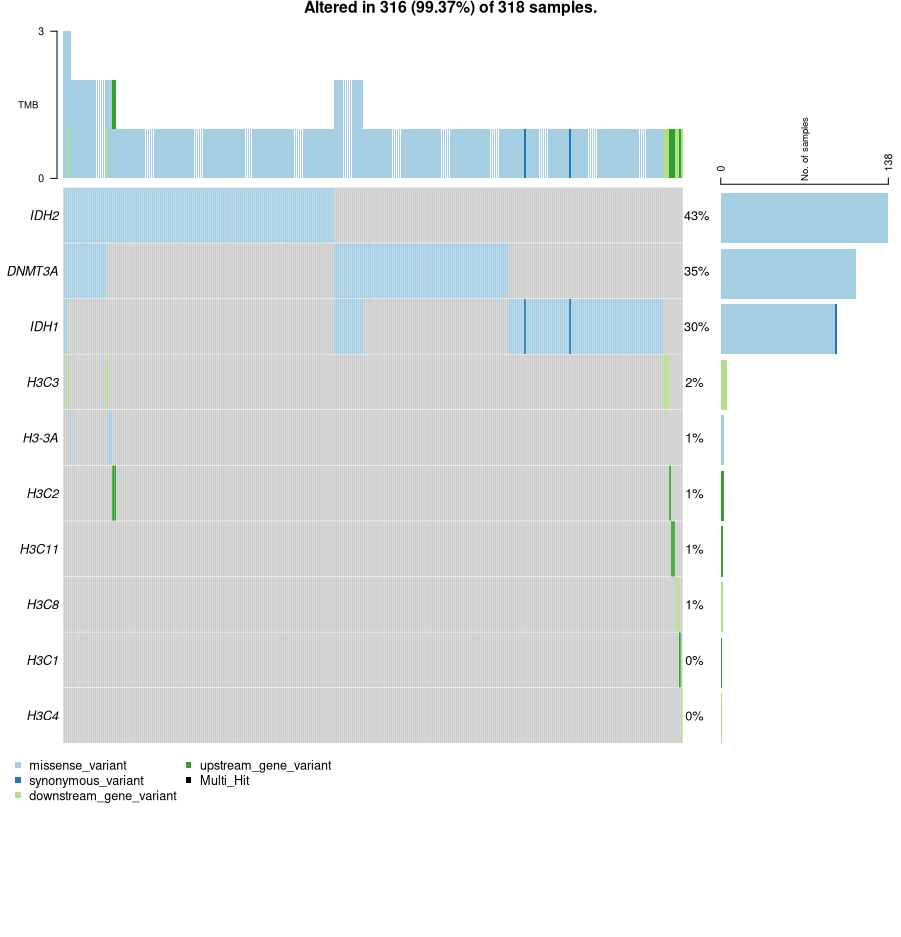

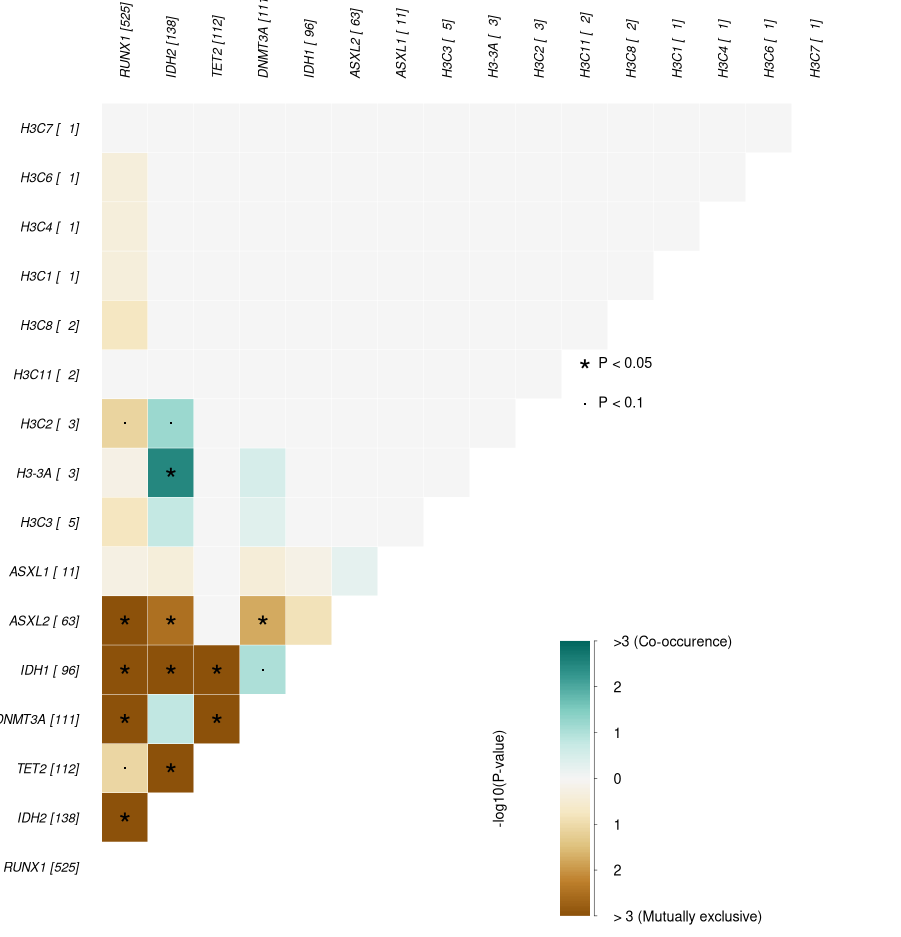

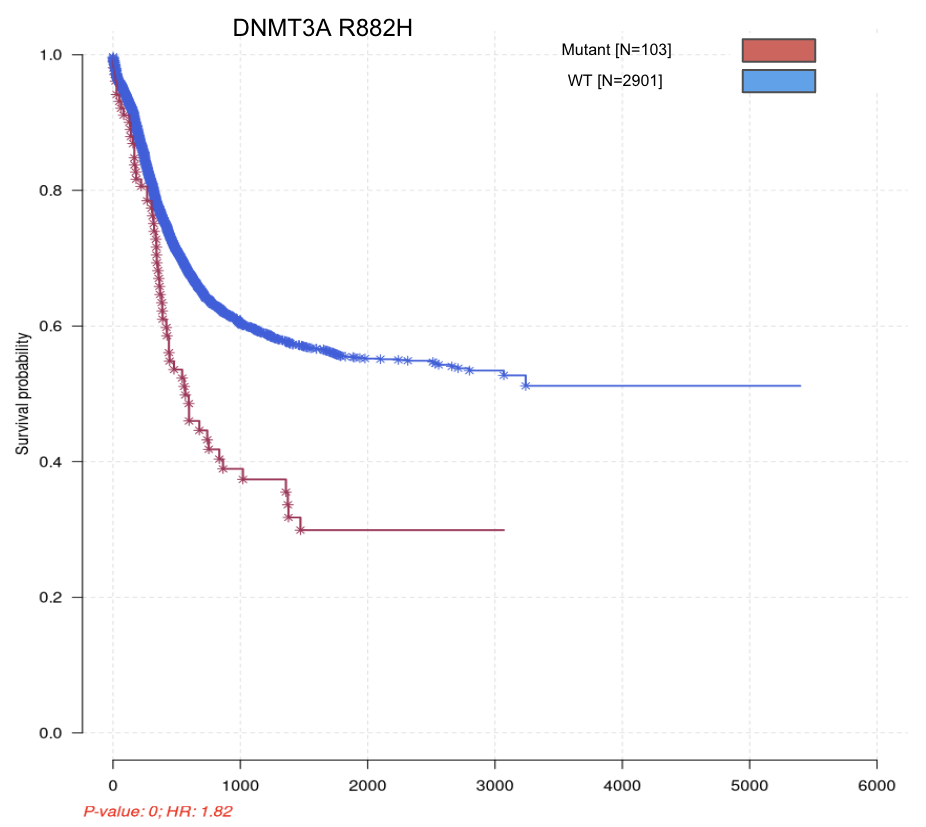

A.

B.

C.

D.

#### Figure S2

**VAF distribution of DNMT3A variants.**

#### Figure S2

**VAF distribution of DNMT3A variants.**

#### Figure S2

### Figure S2

#### **VAF distribution of DNMT3A variants.**

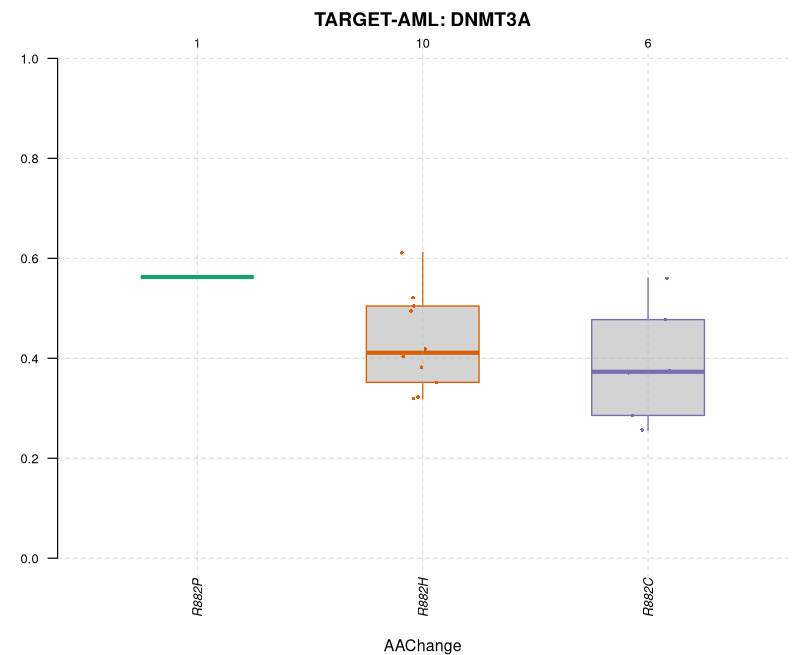

### Figure S3

#### **VAF distribution of H3K27 variants.**

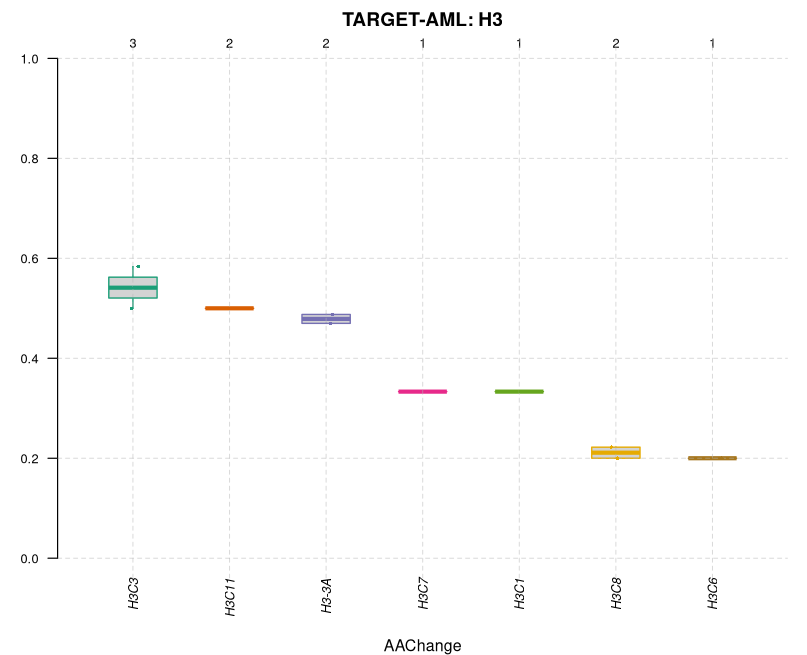

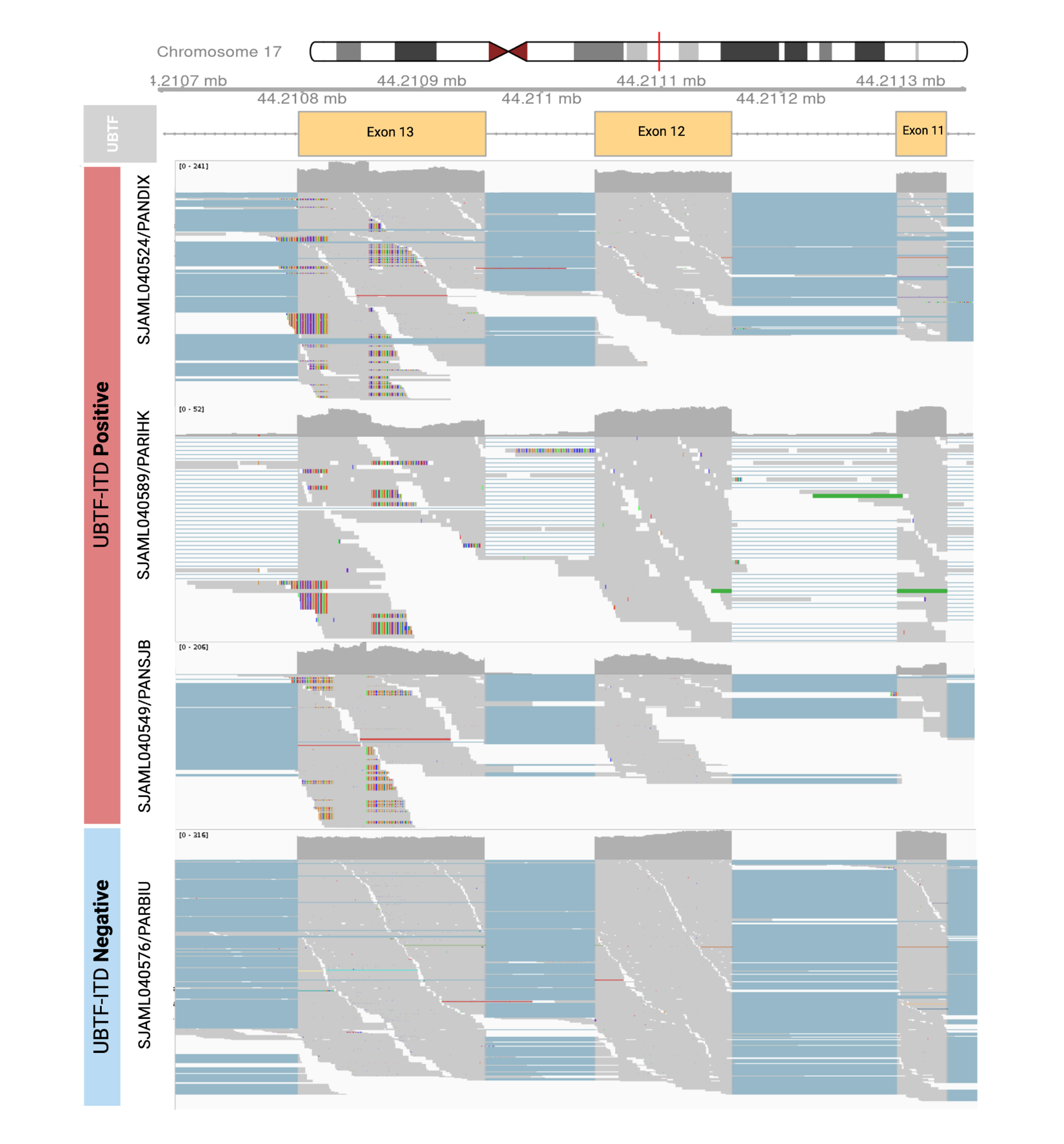

### Figure S4

**Figure S4.** Positive and Negative controls of UBTF-ITD events confirmed in TARGET AML cases. The matched StJude and TARGET IDs of patients are presented. All samples are from primary AML bone marrow. The UBTF-ITD status is confirmed in Umeda, M. et al. (2022) using CICERO(v1.7.0), INDEL detection, and soft-clipped read counts ratio. Integrative Genomics Viewer (IGV) visualization showing soft-clipped reads and increased coverage in UBTF exon 13 in UBTF-ITD positive patients but not UBTF-ITD negative patients.
